# Genomic insight into the biogeographic history, divergence, and adaptive potential of *G. purpurascens*, a forgotten landrace of *G. hirsutum*

**DOI:** 10.1101/2020.09.03.280800

**Authors:** Mian Faisal Nazir, Shoupu He, Haris Ahmed, Zareen Sarfraz, Yinhua Jia, Hongge Li, Gaofei Sun, Muhammad Shahid Iqbal, Zhaoe Pan, Xiongming Du

## Abstract

Continuous selection and adaptation to the local environment resulted in the loss of genetic variation in *Gossypium hirsutum*, which is the most important source of natural fiber. Wild progenitors are an excellent source for strengthening the genetic base and accumulation of desirable traits in modern cultivars. Here we reevaluate a landrace of *Gossypium hirsutum*, formerly known as *Gossypium purpurascens*. We seek to understand the genomic structure, variation, and the adaptive/breeding potential among largely neglected landraces of *Gossypium hirsutum purpurascens*, providing insights into the biogeographic history and genomic changes likely associated with domestication. Population fixation statistics suggested marked differentiation between *G. purpurascens* and current varieties, obsolete accessions, and geographical landraces of upland cotton, emphasizing the divergent behavior of *G. purpurascens*. Phylogeny established the primitive nature of *G. purpurascens*, the inclusion of which in upland cotton gene-pool can enhance the narrowed genetic base of upland cultivars. Genome-wide associations comprehend multiple loci associated with domestication regions corresponding to flowering and fiber quality. Moreover, the conserved nature of *G. purpurascens* can provide insight into understanding the evolutionary process of *G. hirsutum*.

## Introduction

As a natural fiber source, cotton is one of the most important crop in the agriculture industry. Continuously narrowing genetic base has led us to stagnant yield, making it unavoidable to exploit genetic variation as elite alleles present in the distinct gene pool of landraces and wild genotypes (Bolek, El-Zik et al., 2005, Chen, Gols et al., 2015, Tyagi, Gore et al., 2014). Wild progenitors and landraces are excellent sources for developing desirable variation in current cultivars (Huang & Han, 2014). The cotton genus (*Gossypium L*.) consists of 45 diploid species (2n = 2x = 26) and 7 allotetraploid (AD) species (2n = 4x = 52), including *G. hirsutum* L. (AD1), *G. barbadense* L. (AD2), *G. tomentosum* Nuttalex Seemann (AD3), *G. mustelinum* Miersex Watt (AD4), *G. darwinii* Watt (AD5), *G. ekmanianum* (AD6),*G. stephensii* (AD7) (Fryxell, 1992, Gallagher, Grover et al., 2017). Among them, *G. hirsutum* and *G. barbadense* are cultivated species, especially the *G. hirsutum* covered more than 95% cotton-growing areas in the world. *G. hirsutum* has seven geographical landraces, including *yucatanense, punctatum, richmondi, morilli, palmeri, marie*-*galante* and *latifolium*, which have confined distribution in Central America (Wendel, Brubaker et al., 1992). Caribbean islands are considered as the primary center of origin for the wild type of *G. hirsutum* and further expanded along the Antilles chain of islands following the gulf stream, from the coast of Venezuela to the Yucatan and the Florida Keys (Stephens, 1966). The presence of a wild form of *G. hirsutum* on the western coast of Central America can be explained as human-mediated dispersal. Furthermore, these landraces widely dispersed through Pacific ocean currents, on Pacific islands, including Socorro, Wake, Tahiti, Marquesas, Samoa, Fiji, and Saipan islands (d’Eeckenbrugge & Lacape, 2014, Fosberg, 1959, Fryxell & Moran, 1963, Stephens, 1963, Stephens, 1966). This implied that the wild forms of *G. hirsutum* were widely dispersed in the tropical and subtropical islands of the world.

However, *Gossypium hirsutum purpurascens*, a perennial plant, arguably a landrace of tetraploid cotton *G. hirsutum*, was widely distributed in China, India, Congo, Africa, Egypt, and Brazil during the 17^th^ century (Watt, 1927, Watt, 1907). Watt (Harland, 1940, Watt, 1927) classified *G. purpurascens* as a specie of *Gossypium* genus, while Hutchinson (Hutchinson & Stephens, 1944) grouped it as a landrace of *G. hirsutum*. Afterward, *G. purpurascens* was not included among the seven geographical landraces of *G. hirsutum* by the scientists. Thus, we consider it as a forgotten landrace, a potential source of natural variation. *G. purpurascens* was unable to continue its reproductive phase under natural climatic conditions in Central and North China, thus mainly distributed in Hainan province, and was also introduced to Guangdong, Guangxi, and Fujian province in South China where environmental conditions are favorable. In recent years, Dr. Xiongming Du and his colleagues from the Institute of cotton research (ICR), Anyang, China, found native growing *Gossypium purpurascens* in several Islands in South China, i.e., Sansha islands in Hainan Province and Naozhou island near Zhanjiang in Guangdong province (Figure EV1). It is believed that these landraces have been naturally growing on the islands without human intervention for centuries, developing distinct features, i.e., day-length sensitivity, adaptable to local environmental conditions, and distinct plant architecture (Figure S1). The morphological distinctions and the geographic isolation in these islands emphasize that *G. purpurascens* is a wild form *G. hirsutum*. The source and time of the introduction of these landraces have not been reported. However, the presence of wild *G. hirsutum* species in the Pacific sea (d’Eeckenbrugge & Lacape, 2014, Stephens, 1958, Stephens, 1966) and the direction of equatorial currents suggests that the primary geographic origin of *G. purpurascens* should be the western coast of Central America and dispersed to Pacific islands, south China, Australian, Indian and Africa by ocean currents (Figure EV1). Another possible alternative for dispersal might be human-mediated dispersal through trade routes following the ocean currents. Frequent reports by French explorers (Watt, 1907) suggested that *G. purpurascens* might have been distributed to Asia and Africa by French colonists in the 17^th^ century during the first period of introduction of upland cotton in Asia. The distribution areas of the wild and feral *G. hirsutum* and *G. purpurascens* are overlapped, suggesting consideration of both as the primitive forms of *G. hirsutum*. Thus, it is more valuable to understand the genome structure variation and differentiation among these *Gossypium purpurascens* landraces and the different wild form *G. hirsutum* including the central American landraces.

The exploitation of differentiation in upland cotton has been under study for a long time using conventional techniques, i.e., pedigree breeding (Esbroeck & Bowman, 1998, May, Bowman et al., 1995, Tyagi et al., 2014), morphological and biochemical markers (Wendel et al., 1992) and molecular markers (Kuang, Wei et al., 2016, Yu, Fang et al., 2012). In recent years, advances in genomic studies such as Genome-wide association studies (GWAS) enabled us to understand molecular diversity better and dissect the pattern of divergence in cotton cultivars and landraces. Genome-wide association studies have proved to be a useful tool to analyze the genetic profile of complex traits. It has been widely adopted in wheat (Zhai, Liu et al., 2018), maize (Hufford, 2012, Li, Peng et al., 2013), oil crops (Li, Dossa et al., 2018, Rahaman, Mamidi et al., 2018, Wu, Liang et al., 2019), and legumes(Varshney, Saxena et al., 2017, Zhang, Zhao et al., 2019). With the rapid advancement in single nucleotide polymorphism (SNP) genotyping, 3rd generation sequencing has enabled GWAS to provide a better outcome in polyploid crops for association studies. In cotton, GWAS has been used to dissect genetic mechanisms underlying fiber quality traits (Fang, Wang et al., 2017, Liu, Gong et al., 2018, Ma, He et al., 2018, Su, Li et al., 2016), agronomic traits (Huang, Nie et al., 2017) verticillium wilt (Du, Huang et al., 2018, Gong, Yang et al., 2018). However, despite all technological breakthroughs in genomics, contribution towards finding new genetic resources in crop plants has been limited.

To provide the molecular evidence for genomic differentiation and the relationship between different wild and cultivated forms of *G. hirsutum*, we used *G. purpurascens* cotton from South China sea islands, Central American landraces of *G. hirsutum*, current varieties cultivated in main cotton-growing areas in China, and obsolete varieties grown in South China for more than 50 years without management. We re-sequenced 403 accessions and carried out population stratification and genome-wide analysis study (GWAS) for multiple traits. This study will provide insight into genetic differentiation among geographically segregated landraces of *G. hirsutum* and provide the molecular information for oceanic dispersal of wild/feral forms of *G. hirsutum* and the precious gene pool from the forgotten landrace of *Gossypium hirsutum purpurascens*. The excavated information can be further utilized in breeding to expand the narrowed genetic base of upland cotton globally.

## Materials and Methods

### Plant material and phenotyping

Upland cotton accessions (411 genotypes obtained from the cotton germplasm collections from gene bank of the Cotton Research Institute of the Chinese Academy of Agricultural Sciences (CRI-CAAS), were used for phenotyping. These genotypes comprised four distinct groups, i.e., group 1 belongs to current verities (267) currently being cultivated, group two comprised of 95 genotypes collected from south-west China, group three, which is distinct in nature comprised of 49 genotypes (*G. purpurascens*) collected from Sansha Island in Hainan province, Naozhou island and Techeng island of Zhanjiang in Guangdong province (Table S1). While, group 4 (33) comprises of seven reported geographical landraces of *G. hirsutum* i.e., yucatanese, richmondi, morrilli, Marie-Galante, palmeri, punctatum and latifolium (Group 4 was not included in phenotyping). Two replications were planted in six environments for two consecutive seasons in 2017 and 2018. Agro-ecologically different locations comprised of Shijiazhuang (SJZ) in Hebei Province, Changsha (CS) in Hunan province, Anyang (AY) in Henan Province (Yellow River region), Sanya (SAN) in Hainan province, Alaer and Shihezi in the Xinjiang (XJ) autonomous region (Northwest Inland) (Fig. S2). All standard field management practices were applied, including irrigation, pest management, and fertilization, following the usual local management practices in each test location. The cotton was sown in mid-to late April and harvested in mid-to late October at all locations except for Sanya (SAN) where cotton was sown in November and harvested in March.

In all the test locations, phenotypic traits were recorded following the same scoring standard. We characterized fiber-related traits *viz*. Fiber length (FL), Fiber length uniformity (FLU), Fiber elongation (FE), flowering time (DF) and anther color. Flowering was observed daily to record flowering time, and days to flowering (DF) were calculated from the sowing day to the day that the first flower appeared in 50% of the plants in one plot in each environment. Twenty naturally opened bolls from each accession were hand-harvested to gin the fiber for characterization of fiber quality. All samples were subjected to High-volume instrument (HFT9000) for estimation of quality parameters at the Cotton Quality Testing Center in Anyang, China. Data were collected on the fiber length (FL, mm), Fiber elongation (FE %), and fiber length uniformity (FLU %).

### DNA extraction, sequencing, alignment, and SNP detection

We cultured five seeds of each accession in a growth chamber. After the true leaf stage, DNA was extracted from the seedlings. Plant DNA Mini Kit (Aidlab Biotech) was used to extract total genomic DNA, and 350-bp whole-genome libraries were constructed for each accession according to the manufacturer’s specifications (Illumina). Subsequently, we used the Illumina HiSeq X10 by a commercial service “Novogene” platform to generate 6.45-Tb raw sequences with 150-bp read length. SNP calling was performed with GATK standard pipeline (Genome Analysis Toolkit, version v3.1). Sequencing data for G4; GHL was obtained from published data (Fang, 2017).

### Population stratification

Population stratification was examined using ADMIXTURE (Alexander, Novembre et al., 2009) to study population structure, which utilizes clustering method (model-based) to draw population structure assuming different numbers of clusters (K). A total of 190,545 SNPs without missing genotypes were used.

To perform principal component analysis, EIGENSOFT package was used with an embedded SMARTPCA program (Price, Patterson et al., 2006) using 190,545 SNPs without missing genotypes.

To perform Phylogenetic analysis, SNPs of all genotypes were filtered with minor allele frequency MAF=0.05, followed by the construction of a neighbor-joining tree using maximum likelihood method with SNPhylo software (Lee1, Guo et al., 2014). Further, we constructed the neighbor-joining tree using Dendroscope.

### Population genomics analyses

To identify highly differentiated regions, genome-wide population fixation statistics (*F*st) were calculated (SNP data of 404 genotypes were analyzed, excluding low-Quality data and missing data) using vcftools with a sliding window of 100kb and step size of 20kb (--fst-window-size 100000--fst-window-step 20000). The average *F*st of all sliding window was considered as the value at a whole-genome level among different groups.

Highly diverged regions were selected by merging fragments with a distance of less than 50kb after the initial selection of top 1% π values. To identify the putative genes which are under selective pressure between landraces and cultivars, the non-synonymous SNPs with the top 1% of *F*st value were selected, and the corresponding genes were then determined.

Nucleotide diversity (π) is an estimate of the degree of variability within population and species (Tajima, 1983). Nucleotide diversity was calculated using vcftools with sliding window 100kb, based on genotypes in different groups separately.

The reduction of diversity (ROD)/diversity ratios (π_landraces_/ π_cultivars_) between different groups was calculated as an estimate of selection sweeps between different groups. The threshold of top 1% ROD was used to identify significant selection/improvement signals. To further confirm these selection sweeps, the Composite likelihood ratio (CLR) was calculated using SweeD with a grid size of 2000.

### GWAS Analysis

A total of 2,455,716 high-quality SNPs (MAF>0.05, missing rate<20%) in 372 *G. hirsutum* accessions in three sets (Set 1; comprised of 169 accessions whose phenotypic data was collected from SJZ, CS, AY, XJ. Set 2: comprised of 324 accessions whose phenotypic data was collected from AY. Set 3; comprised of 268 accessions whose phenotypic data was collected from SAN. (Accessions with missing SNPs data were excluded from analyses) were used to perform GWAS for multiple traits in efficient mixed-model association expedited (EMMAX) software (Kang, 2010), which was intended to efficiently execute analysis for large-dataset (Li, Yeung et al., 2012). Population stratification and hidden relatedness were modeled with a kinship (K) matrix in the emmax-kin-intel package of EMMAX. The genome-wide significance thresholds of all tested traits were evaluated with the formula P= 0.05/n (where n is the effective number of independent SNPs).

### Statistical analyses

Student’s two-tailed t-tests and one-way ANOVA test were performed in GraphPad Prism software.

## Results

### Population stratification of *Gossypium hirsutum purpurascens*

To scale the genetic differentiation and phylogeny, we performed Population structure, phylogenetic relationship, and principal component analyses using whole-genome SNP data (Figure 1). Phylogenetic analysis, performed by calculating the genetic distance between SNPs, clustered genotypes into four groups (Figure 1c), i.e. current varieties (CV as G1), obsolete varieties from South-West China (SWV as G2), *Gossypium hirsutum purpurascens* landraces (GHP as G3) collected from islands in South China and *Gossypium hirsutum* landraces from Central America (GHL as G4), as supported by principal component analysis (Figure 1b). Groups CV, SWV, GHP, and GHL comprised 215, 89, 43, and 23 genotypes, respectively (Table S2), while 33 lines were found to be admixed. We estimated genome-wide diversity for each group separately. Overall diversity (π) was estimated to be 0.411×10^−3^ for all genotypes while 0.211×10^− 4^, 0.271×10^−4^, 0.244×10^−4^ and 0.453×10^−4^ was observed in group 1, group 2, group 3 and group 4 respectively (Figure 1a). Genetic diversity in modern upland cotton cultivars was significantly lower than landraces, suggesting a stronger domestication bottleneck. Detailed chromosome-wise nucleotide diversity is depicted in Figure S3 and Table S3. GHL showed medium to high diversity within chromosome, while GHP showed lower nucleotide diversity with chromosomes. Although, *G. purpurascens* (group 3) showed relatively less diversity as compared to other landraces, since this group only consist of landraces belongs to the conserved *G. purpurascens*, while group 4 (GHL) includes all seven previously reported landraces. However, the cultivated group showed reduced nucleotide diversity when compared with landraces. The population differentiation statistics (*F*st) between group 1 (CV) and group 2 (SWV) was lower at 0.10647 than other combinations, i.e., GHP vs CV, GHP vs SWV, and GHP vs GHL. *G. purpurascens* showed strong population differentiation compared with CV, SWV, and GHL with *F*st values 0.58103, 0.55377 and 0.4871 respectively (Figure 1a), whereas *G. purpurascens* also significantly differentiated from other landraces belonging to *G. hirsutum viz*. yucatanese, richmondi, morrilli, Marie-Galante, palmeri, punctatum, and latifolium. Earlier reported statistics for genetic diversity and population differentiation are much lower (Du et al., 2018, Fang et al., 2017, Wang, Tu et al., 2017) as compared to our study. Therefore, this implies that the diversity and differentiation among *G. hirsutum* accessions and landraces in this study were higher. Detailed Fst statistics between different groups has been elucidated in Figure S4 and Table S3. It is evident from the provided statistics that GHP showed marked differentiation with other groups when compared on the chromosome level. Genetic differentiation between GHP and other groups ranged from medium to high. Chromosome A05, A06, A07, A08, A10, A11, A13, D01, D03, D04, D06, D07, and D10 depicted higher genetic differentiation as compared to other chromosomes. However, there was a sharp reduction in genetic differentiation when cultivated group and SWV were compared.

**Figure 1:**
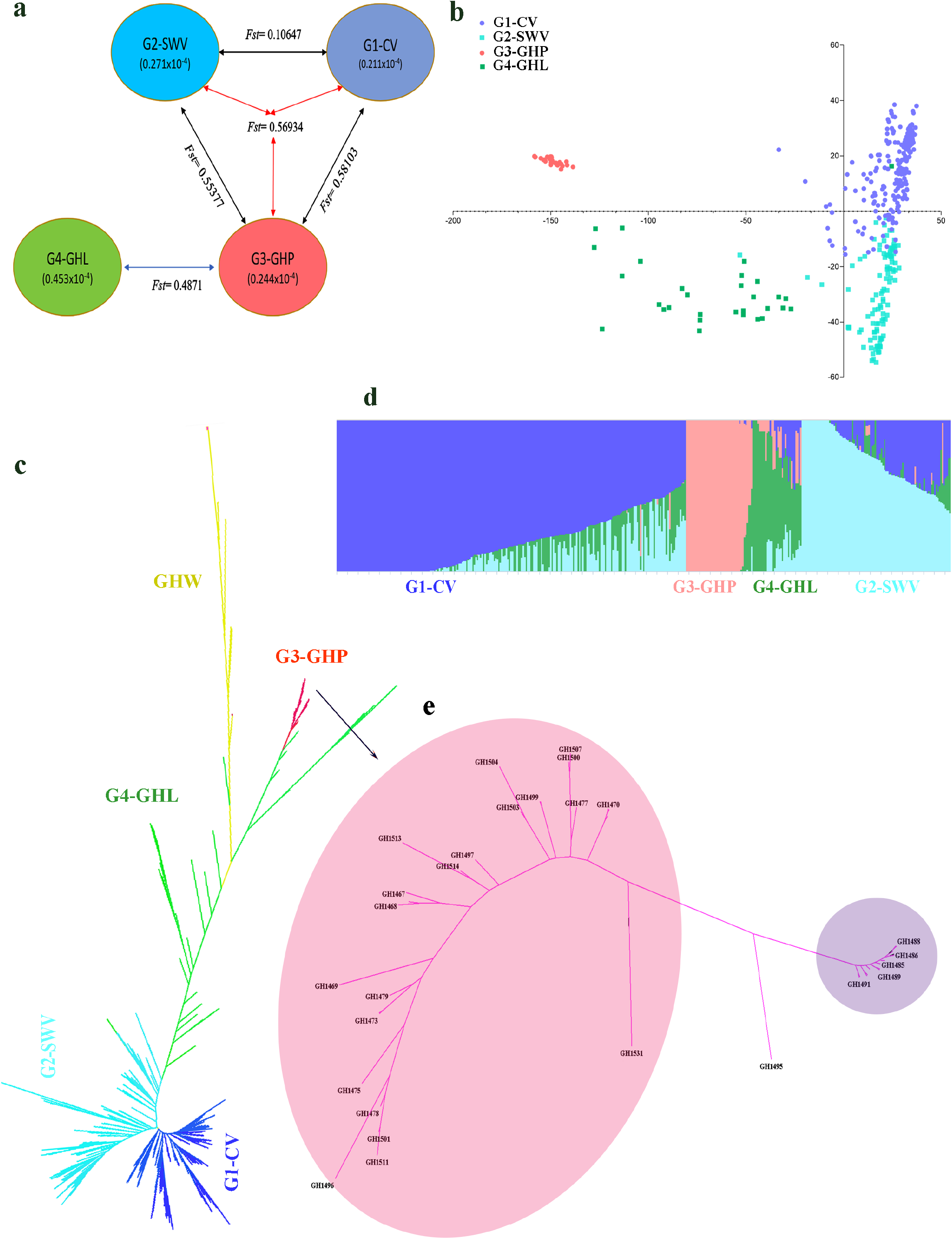
Population Differentiation. **a)** Nucleotide diversity (π) and population fixation index (*F*st) across the three groups. The value in each circle represents a measure of nucleotide diversity for the group, and the value on each line indicates population divergence between two corresponding groups. **b)** Principal component analysis plot of the first two PCAs, i.e., PCA1 and PCA2 for all accessions. Dot color scheme is as Group 1: current cultivars Group2: accessions collected from south china, group 3: landraces collected from Hainan and Guangdong province & Group 4: *G. hirsutum* landraces **c)** Phylogenetic tree constructed using whole-genome data, distributing genotypes into four clades as per original classification. Landraces from Hainan island clustered together in one group **d**) Model-based clustering analysis of all genotypes, performed using ADMIXTURE, representing four groups and their corresponding accessions at K=4 **e)** Phylogenetic tree representing two subclades within *G. purpurascens* (G3-GHP). Purple color represents *G. purpurascens* landraces collected from Sansha island in south China, while light red color represents landraces collected from coastal areas of Guangdong province in south China

Phylogenetic results indicated that *G. purpurascens* was more primitive compared to landraces from Central America except for yucatanese. The phylogenetic tree was constructed using wild relatives of *G. hirsutum* viz. *G. darwinii, G. raimondii, G. ekmanianum*, and *G. mustelinumas* as an outgroup. Detailed phylogenetic results have been presented in Figure 1c, which depicted that the *G. purpurascens* (GHP) and the landraces from Central America (GHL) could be clustered in one clade with two subgroups. *G. purpurascens* accessions were clustered at one end of the clade, and most of the landraces from Central America were in another. *G. purpurascens* showed a close phylogenetic relationship with primitive *G. hirsutum* landraces *yucatanese*, and then *punctatum, Palmeri, Marie-Galante, and latifolium*, finally mixed with other landraces from Central America. This implied that *G. purpurascens* was more primitive compared to landraces from Central America (Fig 1c & Fig. S5). Structure results, in line with phylogenetic analyses, indicated that GHP group is very conserved, even at level K=5. Further, with less heterozygosity, it’s not showing any further differentiation within the group (Figure 1d, Table S2, and Fig. S6).

We further found the population differentiation of *G. purpurascens* was closely related the geographic isolation by the islands. Based on phylogeny, we classified *G. purpurascens* into two subgroups (Figure 1e), which are consistent with the geographic locations of these landraces. Subgroup 1 (indicated with purple color) comprised of landraces collected from Sansha island of Hainan province, while sub-group 2 (indicated with light red color) comprised of landraces collected from Naozhou island and Te Cheng island of Zhan Jiang (coastal areas of Guangdong province in South China). Moreover, population differentiation and reduction of diversity (Fig. S7) among these two subgroups have suggested that chromosomes A02, A06, A07, A10, D02, D05, D08, D09, D10, and D12 have undergone genomic changes due to different geographic distribution in coastal areas of Hainan and Guangdong. Unlike other populations, *G. purpurascens*, having distinct characters, suggested that these genotypes may have a different source of origin (Figure S8 and Table S4). This landrace has been grown in several islands of Hainan and Guangdong province in China, without any human intervention for hundreds of years. Thus, one can argue that this is a purest, naturally acclimatized, preserved, and primitive landrace of *Gossypium hirsutum*, a forgotten source of variation.

### Genetic differentiation between *G. purpurascens* and cultivated genotypes

Genetic differentiation analysis revealed the genome-wide differentiation among the cultivated group and *Gossypium purpurascens*. Pairwise *F*st estimates (Figure S9) presented genome-wide differentiation pattern between *Gossypium purpurascens* (GHP) and the cultivar groups (SWV and CV), which clearly illustrated a decrease in differentiation from GHL to SWV to CV. *F*st estimates for SWV vs CV are much lower when compared with either combination of GHP. GHP showed a high degree of differentiation with CV and SWV throughout the genome except for some regions on chromosome A04, and A12 showed marginalized differentiation among groups, suggested the conserved nature of these regions. The decrease in population differentiation between groups suggests increased selection pressure during the past few decades in developing new high yielding cultivars. To further exploit the genetic divergence within the groups, we combined group CV and SWV (due to low genetic diversity between these two groups), considered as the cultivated group and compared with Gossypium purpurascens (group 3) (Figure 2a and 2c). Fst estimates between the cultivated group and Gossypium purpurascens (Figure 2b and 2d) revealed the presence of genome-wide differentiation among these two groups except for chromosome A04 and A12, which showed very low differentiation. We further narrowed down genomic regions with the highest divergence by selecting the top 1% genome-wide threshold of Fst estimates. We identified 303 regions that harbored 1063 differential genes with 230 non-synonymous and 73 synonymous regions (Table S5). Genes located in these highly diverged regions could potentially play a critical role in shaping modern cultivars.

**Figure 2:**
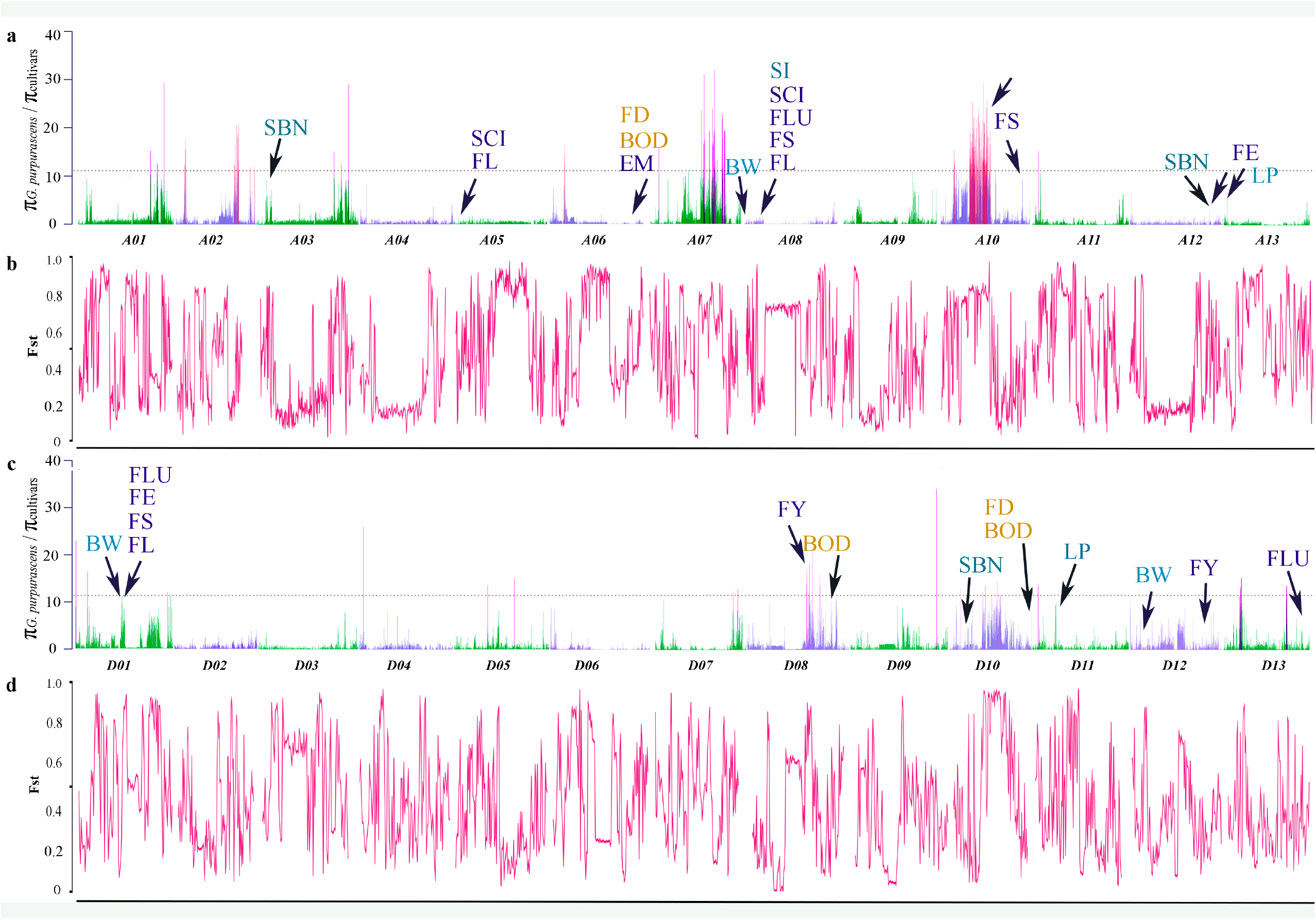
Genetic differentiation between G. *purpurascens* and cultivated genotypes. **a)** π_*G. purpurascens*_/π_Cultivated_ values (genetic diversity in the cultivated group as compare to *G. purpurascens*) for A genome (Chr A01-A13). Group 1 and Group 2 from figure 1 were combined, and π ratio was calculated using whole-genome data with a 100kb sliding window. The horizontal dotted line represents a threshold of top 1% values, whereas the threshold is represented with red columns **b)** Population divergence (*F*st) for A sub-genome (Chr A01-A13). A threshold of the top 1% is selected as highly differentiated regions **c)** π_G. purpurascens_/π_cultivated_ values (genetic diversity in the cultivated group as compare to landraces) for D sub-genome (Chr A01-A13). Group 1 and Group 2 from figure 1 were combined, and π ratio was calculated using whole-genome data with a 100kb sliding window. The horizontal dotted line represents the threshold of top 1% values, whereas the threshold is represented with red columns **d)** Population divergence (*F*st) for D sub-genome (Chr D01-D13). A threshold of the top 1% is selected as highly differentiated regions. \***a**&**b**) Text in blue color represents previously identified QTL hotspots within selection sweeps for Fiber quality(FL=fiber length, FS=fiber strength, FLU=fiber length uniformity, SCI=, the yellow text represents QTLs identified for maturity (FD= flowering days, BOD=days to boll opening), while the turquoise color represents QTLs identified for yield traits (BW=boll weight, SBN=sympodial branch number, LP=lint percentage).

The nucleotide diversity comparison between *Gossypium purpurascens* and cultivar group indicated significant selection sweeps on different chromosomes. Domestication has resulted in many morpho-physiological changes in cotton over the centuries, including annual growth habit, day length sensitivity, seed dormancy, and high-quality fiber. The sharp reduction of diversity (ROD=π_*G. purpurascens*_/π_Cultivated_) is an effective indicator of selection sweeps. To identify selection sweeps, we scanned genomic regions with a marked decrease in nucleotide diversity (π) by comparing *G. purpurascens* and cultivated group. We selected the top 1% reduction of diversity (ROD) values as a threshold, represented with red columns in Figure 2a and 2d. Compared with the cultivar population, we found significant selection sweep regions (total 199 regions genome-wide) on chromosome A07, A10, D08, and D10 in *Gossypium purpurascens* group, which were resulted from human-mediated artificial selection (Figure 2a, 2c, and Table S6). 165 of these identified selection sweeps were present on A-genome, and 34 were present on D-genome. The average ROD for significant selection sweeps in the A-genome (π_*G. purpurascens*_/π_Cultivated_=15.773) was close to the D-genome (π_*G. purpurascens*_/π_Cultivated_=15.061). However, these selective sweep regions between both sub-genomes were largely different. 246 genes were identified to be present in these selective sweeps. Genes present in these selective sweep regions are potentially associated with domestication traits, i.e., seed dormancy, fiber development etc. For instance, RKP (E3 ubiquitin-protein ligase RKP) gene and ERF062 (Ethylene-responsive transcription factor ERF062) gene present in the selective sweep region on chromosome A10 are known for their potential role in positive regulation of seed dormancy (Hu, Zhao et al., 2008, Nonogaki, 2014). Furthermore, among these genes, 18 transcription factors were identified (Table S6), including GRAS (Zhang, Liu et al., 2018), ERF (Long, Yang et al., 2019), HSF (Guo, Liu et al., 2016), WRKY (Li, He et al., 2014, Xu, Wang et al., 2004), bHLH (Yan, Liu et al., 2015), NAC (Zhang, Huang et al., 2018), bZIP (Wang, Lu et al., 2020), NF-YA (Luan, Xu et al., 2014), MYB (Salih, Gong et al., 2016), TALE (Ma, Wang et al., 2019), RAV (Li, Li et al., 2015), and C2H2 (Salih, Odongo et al., 2019). The presence of the above-mentioned transcription factors in regions associated with domestication and selection emphasized the key role of these transcription factors in developing modern cultivars with higher fiber quality and enhanced response towards biotic and abiotic stresses. To further understand the genetic basis of these domestication sweeps, we mapped these selection sweeps with previously known QTLs. Some of the selection sweeps overlapped with previously reported QTL hotspots for multiple traits, i.e., for fiber quality (Liu, Zhang et al., 2016, Liu, Teng et al., 2017, Tang, Teng et al., 2015, Wang, Guo et al., 2006, Wang, Zhu et al., 2012), maturity (Ding, Ye et al., 2015, Liu et al., 2016, Niu, Cai et al., 2018), yield traits (Liu et al., 2017). Whole-genome duplication (polyploidy) provides the raw material for adaption (Hollister, 2015). In the case of *G hirsutum*, D-genome has a significant association of QTLs related to adaption i.e., BOD, DF, and FY, while A-genome has comparatively more associations for traits related to fiber quality. These results are in line with the A and D genome evolution (Xu, Yu et al., 2015).

Furthermore, highly differentiated regions between these two groups overlapped with selection sweeps (Figure 2). This overlapping pattern suggests that this genetic differentiation might have resulted from a rigorous selection over the past century.

### Genetic differentiation between *G. purpurascens* and *G. hirsutum* landraces (Central American)

The significant divergence and selection sweeps were observed when *G. purpurascens* compared with landraces from Central America. Since *G. purpurascens* were dispersed from Central America by ocean currents, we inspected the differentiation of these landraces with the *G. hirsutum* landraces from Central America. These selection sweeps (π_*G. purpurascens*_/π_GHL_) should have resulted from natural selection, so these were not consistent with the human-mediated current selection signals (π_*G. purpurascens*_/π_Cultivated_) (Figure 2a&c and 3a&c) except chromosome A10,which showed both artificial and natural selection signals. This suggested a divergent pattern of selection for modern cultivars. Furthermore, we estimated significantly higher ROD values for improvement (Genome-wide average π_*G. purpurascens*_/π_Cultivated_=1.022) as compare to domestication (Genome-wide average π_*G. purpurascens*_/π_GHL_=0.581), suggesting a stronger selection pressure during the development of modern cultivars. These results are consistent with a narrow domestication bottleneck and stronger improvement bottleneck in cotton improvement.

Population fixation statistics (*F*st) suggested a significant variation on the genomic level between *G. hirsutum* purpurascens and *G. hirsutu*m landraces (Figure 3). Chromosomes A01, A05, A06, A07, A09, A11, and A13 showed a high degree of differentiation on A-genome (Figure 3b) while D-genome (Figure 3d) showed genome-wide differentiation except for chromosomes D12 and D13, which showed comparatively lower differentiation. Furthermore, reduction of diversity indicated some highly significant selection sweeps on chromosomes A03, A06, A10, D02,D04, D07, D12, and D13 when compared *Gossypium purpurascens* with *G. hirsutum* landraces from Central America (Figure 3a and 3c). Above mentioned results for genomic differentiation implicated the divergent nature of *G. purpurascens*, which has been the result of natural selection in an isolated area over the centuries. Our results also emphasized on the conserved and primitive nature of *G. purpurascens* in population structure. We propose that the landrace *Gossypium hirsutum purpurascens* should be considered as an important and primitive source of natural variation; thus it can be included with the other seven geographical landraces of *G. hirsutum* from Central America i.e., Yucatanese, richmondi, morrilli, Marie-Galante, palmeri, punctatum, and latifolium.

**Figure 3:**
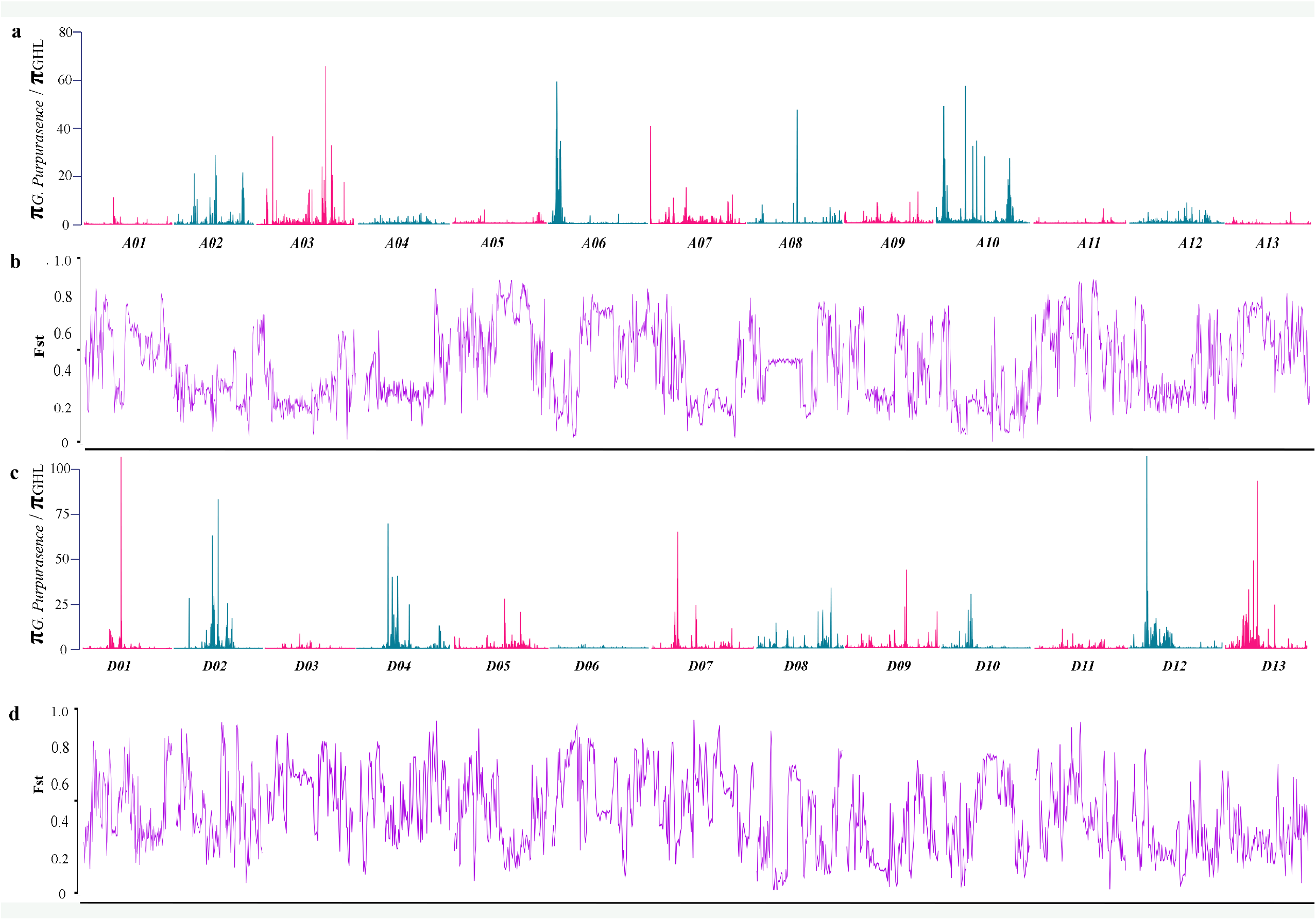
Genetic differentiation among *G. purpurascens* and other *G. hirsutum* landraces. **a)** π_*G. purpurascens*_/π_GHL_ values (genetic diversity in G. *purpurascens* (group 3) as compared to *G. hirsutum* landraces (group 4)) for A sub-genome (Chr A01-A13) **b)** Population divergence (*F*st) for A sub-genome (Chr A01-A13). **c)** π_*G. purpurascens*_/π_GHL_ values (genetic diversity in *G. purpurascens* (group 3) as compared to *G. hirsutum* landraces (group 4)) for D sub-genome (Chr A01-A13) **d)** Population divergence (*F*st) for D sub-genome (Chr D01-D13)

### Genome Wide Association Study for flowering time

Ever-increasing industrial pressure and improvement of cultivars resulted in narrowed genetic diversity in the gene pool due to rigorous selection for fiber yield, fiber quality, early maturity, and seed dormancy. To understand the causal elements behind changes in adaptive traits, we performed GWAS for flowering time, fiber yield, and fiber quality. Association results identified 264 significant associations for flowering time, fiber yield, and fiber quality, 91 of which were either present in exonic/intronic or upstream/downstream of the genes present in the region (Figure S10 and Table S7). Five significant GWAS signals for the flowering time were identified to overlap with the selection sweep and high differentiation regions on chromosome A05, A06, D01, D02, and D10 (Figure 4 and Fig. S11 and S12). QTL region 19.0Mb-19.3Mb (Figure 4g) on chromosome A05 corresponds to 13 genes, 30Mb-70Mb (Figure 5c & f) on chromosome A06 (Figure S12) corresponds to 217 genes among which 51 non-synonymous SNPs were identified, and the QTL region on Chromosome D01 22.37Mb-24.72Mb (Figure S12) corresponds to 36 non-synonymous SNPs (Table S7). We identified three genes on chromosome A05, which are related to the flowering phase. QWRF2 (QWRF motif-containing protein 2) involved in phase change, vegetative to the flowering stage (Xu, Jiang et al., 2016). LBD39 (Lateral organ body (LOB) domain-containing protein 39) involved in key coordinated role in the expression of nitrogen assimilation genes and the expression of nitrate reductase (Albinskya, Kusanoa et al., 2014, Yanagisawa, 2014, ZHANG, ZHANG et al., 2014), moreover expression of nitrate reductase genes is influenced by day length, low temperature, and drought, which influence flowering time in plants (Tanaka & Takimoto, 1978). APT1 (Adenine phosphoribosyl transferase) causes male sterility in plants (Gaillard, Moatt et al., 1998, Zhang, Guinel et al., 2002). Expression profiles of the above-mentioned genes have been presented in Figure S13. Genotyping for causal SNPs for the flowering time indicated the transformation of locus GG (haplotype A) to AA (haplotype B) was resulted, either from domestication or human selection. Haplotype GG (corresponds to *G. purpurascens*) shared 8.14% and 7.06% variation on chromosome A05 and A06, respectively.

**Figure 4:**
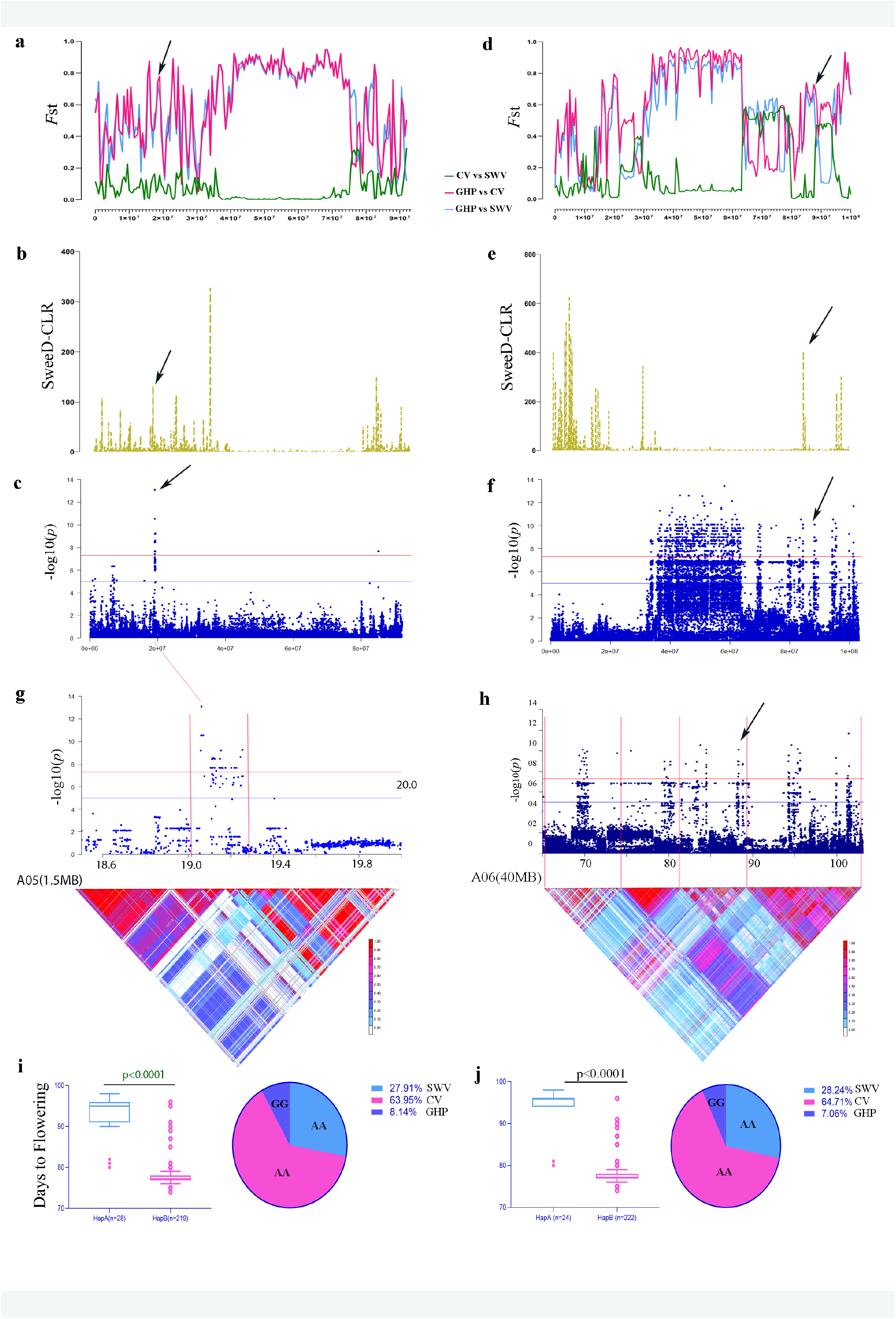
GWAS for the flowering time. **a)** Pairwise *F*st estimates *viz*. G1vsGa, G1vsG3, and G2vsG3 for chromosome A05 **b)** SweeD-CLR score for chromosome A05, peaks indicates the selection sweeps **c)** Local Manhattan plot of chromosome A05 for days to flowering, peak indicates significant GWAS association and Red horizontal line indicates the genome-wide significance threshold **d)** Pairwise *F*st estimates *viz*. G1vsG2 (CV vs SWV), G3vsG1 (GHP vs CV) and G3vsG2 (GHP vs SWV) for chromosome A06 **e)** SweeD-CLR score for chromosome A05, peaks indicates the selection signals **f)** Local Manhattan plot of chromosome A06 for days to flowering, peak indicates significant GWAS association **g)** Expansion of candidate region (18.5mb-20mb) (Top) & LD heatmap for of corresponding region (Bottom) **h)** Expansion of candidate region (30mb-70mb) (Top) & LD heat map for of corresponding region (Bottom) **i)** Box and whisker plot for days to 50%flowering for the two haplotypes where n=28 &219 for subsequent haplotypes, while circle on the right side presents percentage variation shared by each group. SWV and current varieties sharing the same allele corresponds to 91.86% of the variation while GHP (*Gossypium purpurascens*) corresponds to 8.14% of variation **j)** Box and whisker plot for days to 50%flowering for the two haplotypes where n=24 &222 for subsequent haplotypes, while circle on the right side presents percentage variation shared by each group. SWV and current varieties sharing the same allele correspond to 92.95% of the variation, while GP (*Gossypium purpurascens*) corresponds to 7.06% of the variation.

Chromosome A06 showed a significant association with flowering time consistent with the previous report (He, Sun et al., 2018). In addition to previously reported QTLs, we identified more QTLs hotspot on chromosome A06 (Figure 4h and Figure S11). To identify putative candidate genes associated with flowering time, we further determined the LD blocks in the neighboring regions of the significant loci (-log_10_ (*p*) > 6). Five LD blocks on chromosome A06 were exploited (Figure 4f & h and Figure S11). Significant GWAS loci on chromosome A06 (A06_88332071) was present in a highly differentiated region, as depicted in Figure 4d. Furthermore, this locus was also confirmed to be underlying selection with CLR (composite likelihood ratio). Haplotype analysis for A06_88332071 showed a differential pattern for *G. purpurascens* and cultivated group. Haplotype A with GG alleles corresponds to *G. purpurascens* (covering 7.06% of the variation). In comparison, haplotype B with alternate alleles AA corresponds to the cultivated group (covering 92.94% variation with 64.71% share of current varieties and 28.24% share of Southwest varieties) (Figure 4j). this differential pattern of allelic distribution suggests that this locus might have been transformed due to domestication or selection. The other significant associations for flowering time (present in 60Mb-105Mb region and 30Mb-40Mb region) were also mapped for neighboring LD blocks (Figure 4h and Figure S11).

Another Significant GWAS association for the flowering time was identified on chromosome D01 (Figure S12). We determined LD blocks corresponding to the significant loci (Figure S12). This locus was present in a highly differentiated region (identified using *F*st statistics). Furthermore, the genotyping analysis revealed two haplotypes associated with early and late flowering. A list of putative genes identified as a result of GWAS associations has been presented in Table S8. A total of 105 putative genes were identified for flowering time. Further exploitation of these genes can provide useful information to understand the flowering mechanism in cotton.

### GWAS for fiber quality

We identified significant GWAS signal on chromosome D11 for fiber quality traits, i.e., Fiber length (FL), fiber length uniformity (FLU), and fiber elongation (FE) on a region significantly differentiated among populations (Figure S14). We identified 62 key SNPs for fiber quality, but we only focused on pleiotropic loci for FL, FLU, and FE, as these traits are positively correlated to each other. Our results suggested that this region has been under selection for fiber quality. In this region, we found two linkage disequilibrium blocks significantly associated with SNPs. Block 1 (Figure S14 h and i) contain three genes viz. CYCP 3, DGP3, and PLP9 and block 2 (Figure S14 i) contains two genes viz. bZIP 25 and Gh_D11G1318. We generated two haplotypes for each trait based on identified key SNPs. Haplotype A (n=13) consists of *G. Purpurascens* (Group 1) with allele GG, while haplotype B consist of CV (Group 1) and SWV (Group 2) with allele AA. Interestingly haplotype A was associated with relatively higher fiber quality, which is also depicted by average statistics for fiber quality Table S4. Genes present in first LD block CYCP 3; a cell cycle related gene regulated by overexpression of GbTCP (TCP transcription factor in cotton which directly influence fiber elongation and root hair development), might have a secondary role in fiber development (Hao, Tu et al., 2012). DGP3; represents DAR ortholog for chloroplast in higher plants is significantly associated with DAR GTPase activity, which has been characterized for cellular differentiation and morphogenesis of ovule in Arabidopsis (Hill, Broadhvest et al., 2006). While gene present in second LD block bZIP25; a transcription factor which has been involved in seed maturity in Arabidopsis (Alonso, Onate-Sanchez et al., 2009, Santos-Mendoza, Dubreucq et al., 2008), as it is evident that seed maturity is directly correlated with fiber maturity in cotton, thus it can be suggested that this gene might have played an indirect role in fiber development and maturity. Expression profiles of the above-mentioned genes have been presented in figure S15.

## Discussion

*G. purpurascens*, a primitive landrace of *G. hirsutum*, has been naturally acclimatized in an isolated territory (South China sea islands) without any human interventions, arguably having a rich source of unexplored variation, can provide us with tools to manipulate narrowed diversity and avoid the threat of genetic vulnerability in cotton. In contrast with other *Gossypium* species, *Gossypium hirsutum* occupies a relatively wide geographical range. Localized small populations of wild and feral *G. hirsutum* have been found from South America (Venezuela and North-east Brazil) through Caribbean islands, to Central America (Yucatan, El Salvador, and Guatemala) and islands in the Pacific Ocean (Socorro, Wake island, Saipan, Fiji, Samoa, Tahiti, and Marquesas) (d’Eeckenbrugge & Lacape, 2014, Gallagher et al., 2017, Stephens, 1966, Stephens, 1977). The distribution pattern (Watt, 1927) of *G. purpurascens* in central America, South America, Asia, and Africa coincide the direction of ocean currents suggested its transoceanic dispersal from Central America to other parts of the world. Transoceanic dispersal plays a key role in maintaining connectivity among plant populations (Miryeganeh, Takayama et al., 2014). Wild *Gossypium hirsutum* are native to coastal habitats with the potential of transoceanic dispersal as their seeds have a hard seed coat with a tendency to tolerate seawater for a long time without compromising the viability of seeds (Stephens, 1958). Evidence suggested the presence of truly wild *G. hirsutum* species in the North Pacific Ocean (Socorro island, Wake Island, and Saipan island) and South Pacific Ocean (Fiji, Samoa, Tahiti, and Marquesas) (Fryxell, 1965, Fryxell & Moran, 1963). Another example of long-distance transoceanic seed dispersal is the presence of primitive decedents of *G. hirsutum* in Australia (Fryxell, 1966, Fryxell, 1979). The overlapped distribution pattern of geographic landraces of *G. hirsutum purpurascens* and *G. hirsutum* landraces in Central America, i.e., Mesoamerican landraces and Caribbean landraces, suggests *G. hirsutum purpurascens* as a wild form of *G. hirsutum*.

Geographic distribution (confined to coastal areas), morphological characteristics (perennial in nature), and genome-wide differentiation statistics proposed that *G. purpurascens* is also agronomically primitive and is not derived from modern/elite cultivars. We also perceived that *G. purpurascens* was used as a synonym for the wild type *G. hirsutum* after dispersing from the place of origin, i.e., Central America. The wild form *G. hirsutum* was dispersed on the isolated islands (Socorro, Wake island, Saipan, Fiji, Samoa, Tahiti, and Marquesas) in the Pacific Ocean by the ocean currents. In this study, we have found the evidence for the presence of *G. purpurascens* in two islands (Sansha Island and Naozhou Island) in South China, having genome differentiation. This implied that the geographic isolation of the different islands in the Pacific Ocean might have resulted in morphological distinctions and genetic diversity of the wild form cottons on these islands. Therefore, we contend that the cottons (*Gossypium stephensii*) of Wake Atoll island located in the Western Pacific represent a new species of *Gossypium* (Gallagher et al., 2017), and we argue that these are the wild type of *G. hirsutum*. The diversified genomic background of *G. purpurascens* among all the islands along the Ocean currents from Central America to the Pacific and other Ocean islands will provide a rich gene pool for tetraploid cotton. We also believe that the exploitation of all the wild form *G. hirsutum* will make comprehend the genomic evolution in tetraploid *Gossypium*.

Presented results suggested wide genetic differentiation among *G. purpurascens*, modern cultivars, obsolete accessions from South-West China, and other landraces of *G. hirsutum*. Phylogeny, principal component, and structure analyses revealed the divergent behavior of *G. purpurascens* and *G. hirsutum* landraces compared to modern cultivars that were in accordance with genetic differentiation analysis. The divergent trends of landraces compared to modern cultivars were also found in support of previous studies (Fang et al., 2017, Qi, Liu et al., 2013, Varshney et al., 2017, Wang et al., 2017, Xu, Liu et al., 2011, Zhou, Jiang et al., 2015). In contrast with previously reported work, suggesting narrowed genetic diversity among studied cultivars of upland cotton and also in other crops (Rana, Singh et al., 2005, Wang et al., 2017, Wendel et al., 1992), our results pointed out marked differentiation at the genetic level corresponding to *G. purpurascens*. Besides, we identified many selection sweeps and differentiation regions, suggesting the domestication bottlenecks. Genetic bottlenecks associated with crop domestication may have resulted in the loss of genetic diversity and elite alleles. Wild progenitors and landraces are excellent sources for developing desirable variations in current cultivars (Huang & Han, 2014). *G. purpurascens*, a primitive landrace of *G. hirsutum*, has been naturally acclimatized in an isolated territory (South China sea islands) without any human interventions, arguably having a rich source of unexplored variation, can provide us with tools to manipulate narrowed diversity and avoid the threat of genetic vulnerability in cotton germplasm.

Further, to understand the genetic basis of domestication and improvement in flowering and fiber quality, we compared the location of selection sweeps with the significant loci of GWAS analysis. Moreover, loci associated with the flowering time on chromosome A05, A06, and pleiotropic loci associated with fiber quality traits *viz*. FL, FLU, and FE on D11 fall within the selection sweeps. Selection of these loci likely to be involved in fiber quality improvement and implying that during the process of domestication, breeders have selected these regions for improving fiber quality. Compared to previous reports, these loci for fiber quality are also novel (Ma et al., 2018, Wang et al., 2017).

We identified the set of candidate genes that were significantly associated with flowering time and fiber quality traits. The functional characterization of these genes in the future will help us to understand the genetic mechanisms underlying these genes. Our study not only presents a genetic basis to understand the domestication process but also provides a comprehensive insight into the diversification of forgotten source of variation *G. purpurascens* which is necessary for modern-day breeding to utilize in understanding and manipulations of mechanisms for improvement of modern upland cotton cultivars.

## Acknowledgement

We thank Mr. Jueming He from Guangdong Ocean University to provide seeds of *G. purpurascens* collected from islands in the coastal areas of Guangdong province. Special thanks to Prof. Dr. Muhammad Kausar Nawaz (PMAS AAUR) for proofreading the manuscript.

## Data Access

The datasets used and analyzed during the current study are included in this published article and its supplementary information files. All the DNA sequence data have been deposited into the Public database of National Center of Biotechnology Information (NCBI) under accession number PRJNA605345.

## Authors’ contributions

XD and MFN conceived and designed the research project. MFN, SH, and GS performed the main genotyping and other bioinformatics analyses. HA, ZS, and MSI assisted in phenotyping. YJ and ZP cultivated all the plant materials and helped in phenotypic data collection. MFN analyzed all the data and wrote the manuscript. XD and SH revised the manuscript. All authors read and approved the final manuscript.

## Figure legends

**Figure EV1: Geographic distribution of truly wild *G. hirsutum* (TWC) (Fryxell, 1966), geographical landraces of *G. hirsutum (d’Eeckenbrugge & Lacape, 2014)*, and dispersal of *G. purpurascens* through ocean current**.

**a)** Map representing dispersal of *G. purpurascens* through equatorial currents in Pacific Ocean. Horizontal red lines indicate the direction of equatorial currents in the Pacific sea. Cotton flowers in the Pacific Ocean represent the islands where wild *G. hirsutum* has been reported (Socorro island, wake island and Saipan island in the North Pacific Ocean while Fiji, Samoa, Tahiti, and Marquesas are in South The Pacific Ocean, while *G. purpurascens* has been reported in the South China Sea, India, Cango, Brazil, and Australia (Watt, 1907) **b)** Detailed geographic distribution of *G. purpurascens* island in South China sea.). The red dotted circles represent islands (Sansha island, Sanya, Hainan island, Techang island, and Naozhou island) in South China, from where *G. purpurascens* were collected. **c)** Represents the geographic distribution of *G. hirsutum* landraces in Caribbean Sea. Red dots represent the geographic distribution of Caribbean landraces while purple dots represent the geographic distribution of Mesoamerican landraces.

## Supplementary Information

Supplementary Table S1. list of accessions, cultivars, and landraces used in this study (Separate file)

Supplementary Table S2. Structure results and corresponding grouping for all accessions and landraces used in this study (Separate file)

Supplementary Table S3. Chromosome wise nucleotide diversity and pairwise estimates of Fst

Supplementary Table S4. Morphological characteristics of CV, SWV, and GHP in five locations

Supplementary Table S5. List of highly differentiated regions and genes located in the highly differentiated regions identifies through pairwise *Fst* estimates (top 1% threshold) (Separate file)

Supplementary Table S6: Selection sweep regions, Top 1% threshold of π_G. *purpurascens*_/π_Cultivated_

Supplementary Table S7. Key SNPs identified for flowering time, fiber quality and anther color, and their corresponding annotations (Separate file)

Supplementary Table S8. Candidate gene identification for Flowering time, fiber quality and anther color (Separate file)

Supplementary Figure S1. A pictorial description of the *G. purpurascens*

Supplementary Figure S2. Geographic distribution of accessions and landraces used in this experiment

Supplementary Figure S3. Nucleotide diversity of three group of accessions including GHL=G. hirsutum Landraces, GHP=G. hirsutum purpurascens, and cultivated group

Supplementary Figure 4. Chromosome-wise Fst estimates between different groups under study GHL=G. hirsutum Landraces, GHP=*G. hirsutum purpurascens*, CV=current varieties, SWV=South west varieties

Supplementary figure S4. Detailed phylogenetic tree for GHP (*G. purpurascens*) and GHL (landraces of *G. hirsutum*).

Supplementary Figure S6. Structure results k2 to K9. The dotted line represents landrace *G. Purpurasens*, depicting the conserved nature of this landrace.

Supplementary Figure S7. Genetic differentiation two sub-clades in *G. purpurascens*

Supplementary Figure S8. Comparison of morphological characteristics of *G purpurascens* Landraces with other geographical landraces of *G. hirsutum*

Supplementary Figure S9. Pairwise estimates of Population fixation statistics (*Fst*) between each group

Supplementary Figure S10. Manhattan plot for GWAS results of a) Days to flowering b) Fiber Length c) Fiber elongation d) Fiber length Uniformity

Supplementary Figure S11. GWAS for flowering and Local Manhattan plot for chromosome A06 (65Mb-108Mb)

Supplementary Figure S12. GWAS for flowering and identification of candidate genes influencing flowering

Supplementary Figure S13. Expression profiles of putative genes identified for involvement in flowering Supplementary Figure S14. GWAS for Fiber quality traits

Supplementary Figure S15. Expression profiles of putative genes identified for involvement in fiber quality

Supplementary Figure S16. GWAS for Anther color

